# Reference-free and *de novo* Identification of Circular RNAs

**DOI:** 10.1101/2020.04.21.050617

**Authors:** Yangmei Qin, Tingting Xu, Wenbo Lin, Qingjie Jia, Qiushun He, Ke Liu, Juan Du, Linshan Chen, Xiaozhen Yang, Fei Du, Mengjun Li, Min Chen, Tao Tao, Zhi-Liang Ji

## Abstract

**Background:** Early studies have unveiled multiple regulatory functions of circular RNAs (circRNAs); however, accurate detection and quantification of circRNAs, especially in understudied organisms, is unsolved yet.

**Results:** In this study, we developed a new reference-free method, namely Cirit, to *de novo* detect circRNAs with sequence support from the next generation sequencing (NGS) transcriptome. The Cirit showed remarkable performance in accurate detection and quantification of circRNAs via comparing with current methods from different aspects. Using Cirit, we detected 28,813 nonredundant human circRNAs from 91 transcriptome datasets, as well as 2,385 circRNAs from four understudied organisms. Subsequent analyses found that the majority of human circRNAs expressed spatiotemporally; only a very small portion of circRNAs were back-splice junction (BSJ)-consistent (maximally 5.17%) or sequence-consistent (under 2%). Furthermore, circRNA genesis were relatively flexible that only about 60% of human circRNAs used the canonical GT-AG pattern for back-splice. The inconsistent expression of circRNAs challenges their roles as precise transcriptional regulators.

**Conclusions:** In summary, the reference-free method provides a straightforward and universal way for reliable circRNA research. It will largely boost the successful rate in designing highly specific probes to monitor circRNA behavior in cell. In particular, it brings circRNA research to the organisms that have no genome or draft genome.

## Background

Circular RNA (circRNA) is a kind of noncoding RNAs (ncRNAs) generated mainly from back-splicing of pre-mRNA, in which the upstream splice acceptor (SA) is spliced to the downstream splice donor (SD) [1-3]. Current knowledge considers circRNA not a transcriptional splice error but undertaking biological functions like microRNA (miRNA) and protein sponge [4, 5]. Several studies also demonstrated that circRNAs could serve as templates for protein translation [6-8], producing active small proteins or peptides [9, 10]. In particular, many studies designed exquisite experiments to depict circRNA as an efficient sponge for miRNA absorption in the cytoplasm, via which circRNA can interfere gene transcription. However, before we move forward to explore circRNA functions or even its potential clinical applications, many questions need clear answers. For instance, have the circRNAs been detected confidently with sequence evidence? Do the circRNAs express consistently? Have the expressions of circRNAs been quantified accurately?

In recent years, a number of algorithms were introduced to detect circRNAs from RNA sequencing (RNA-seq) transcriptome in large scale. These algorithms, including find_circ [4], CIRCexplorer [11], circRNA_finder [12], CIRI [13], KNIFE [14], NCLscan [15], DCC [16], and UROBORUS [17], usually take the back-splice junction (BSJ) spanning reads as the critical clues in circRNA detection [4, 11, 13, 18]. They discover BSJs by either search of the wrong bridging events between the annotated exons (annotation dependent) or identification of potential splice patterns via mapping reads back to the reference genome or pseudo-sequences (annotation independent) [19]. The performance of BSJ discovery highly depends on splice alignment against a reliable genome [20], which will inevitably produce false positives due to the complexity of splice identification and sequence alignment. Although each of these algorithms have implemented a distinct set of heuristics or rules to minimize false positive rate in BSJ discovery [20], recent comparison analyses revealed regretfully that different algorithms often yielded heterogeneous results from the same sequencing transcriptomes [21, 22]. In addition, most algorithms claim circRNA without providing full-length sequence as a direct evidence in advance. Indeed, some methods such as circseq_cup [23] and the oven-fresh CIRC-full method[24] can generate full-length sequence for circRNA however only after the BSJ determination, which the accuracy and integrity of the circRNA sequences are largely not validated yet. Since there is still lack of a benchmark or gold standard to evaluate circRNA detection fairly, only a small portion of circRNAs predicted by the reference-based methods can be experimentally validated [23]. Moreover, accurate detection and quantification of circRNA isoforms for a definite BSJ is far beyond the scope of current methods. Therefore, current knowledge of circRNA, including its sequential characteristics, expression, quantification, biogenesis, and biological functions, could be incomplete or even deflected.

In this study, we aim to break the constraints on current circRNA detection methodologies and introduce a novel reference-free method for genome-wide detection of circRNAs from RNA-sequencing (RNA-seq) transcriptome. In addition, we will re-characterize human circRNAs in both aspects of sequence composition and expression consistency.

## Results

### Performance evaluation of the new algorithm Cirit

#### Large-scale detection of circRNAs by Cirit

In this study, we developed a new reference-free algorithm, namely Cirit, to *de novo* detect circRNAs from RNA-seq transcriptome (Figure 1 & Methods). To test the feasibility of new method, we prepared two parallel transcriptomes for the cultured HepG2 cell line in basis of the circRNA-enriched library (poly(A)^-^/ribo^-^) and the total RNA library (ribo^-^). Respectively, we detected 8,182 and 1,660 sequence-supported circRNAs using the Cirit. The low circRNA detection rate in the total RNA transcriptome is mostly because circRNA signals were substantially overwhelmed by the linear RNAs during sequencing [20]. Of the 8,182 circRNAs detected in the poly(A)^-^/ribo^-^ transcriptome, 4,568 were *de novo* identified with full-length sequence support, and 3,614 were incompletely assembled however possessed an intragenic-fusion structure when mapping the partial sequences back to the human genome (GENECODE, GRCh37.p13). Likewise, of the 1,660 circRNAs detected in the ribo^-^ transcriptome, 463 were *de novo* identified with full-length sequence support, 1,197 were partially assembled. Besides, for the circRNAs identified from both transcriptomes, 7,689 and 1,584 circRNAs can be annotated by referring to the human genome with high confidence. These results manifested that the Cirit could work on transcriptomes determined from both circRNA-enriched and total RNA libraries.

**Figure 1.**
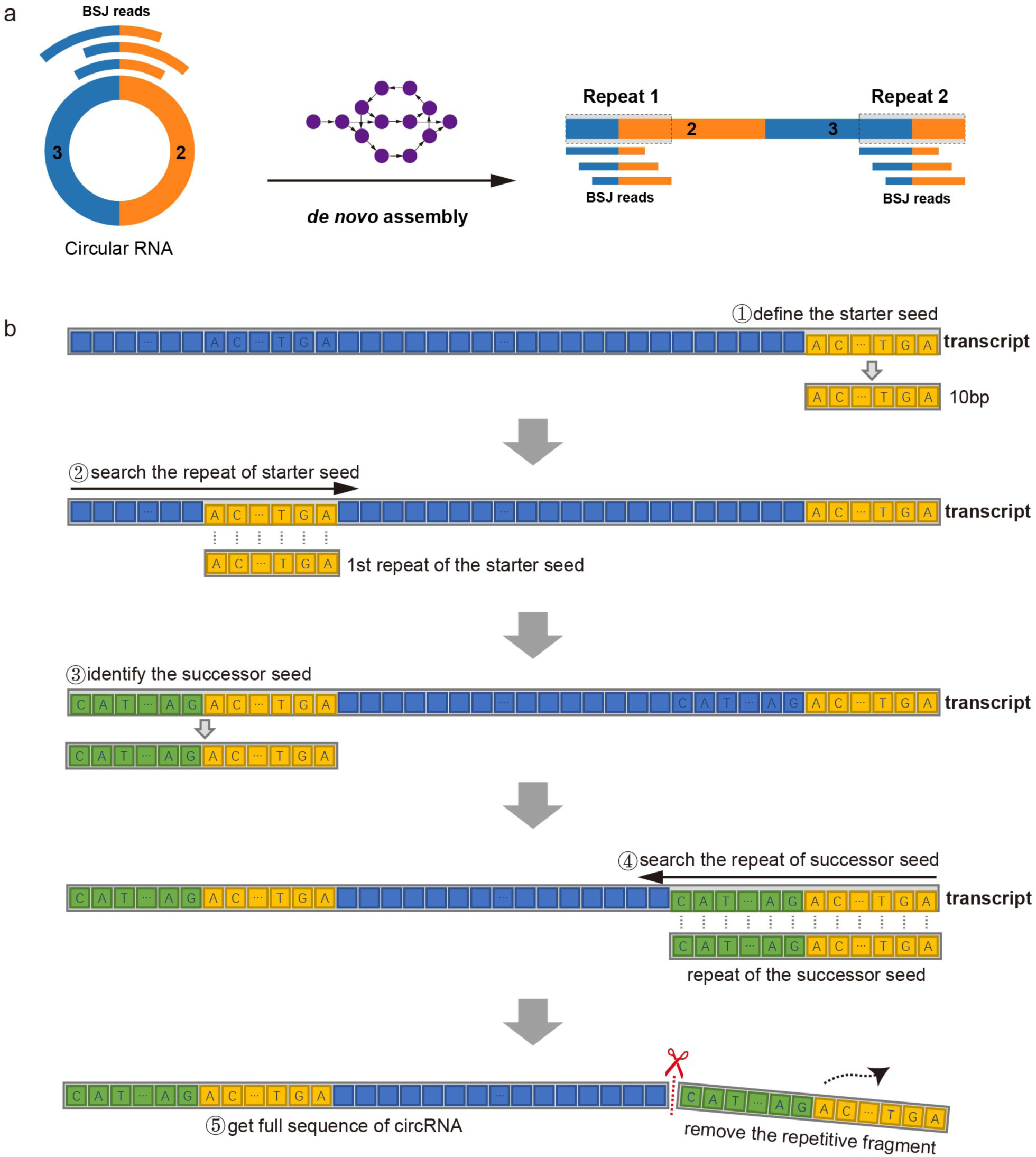
The algorithm of reference-free circRNA detection. **a.** The head-to-tail overlapping structure. When applying *de novo* assembly method to reconstruct circRNAs, the “back-splice junction-spanning reads” will lead circRNA assembled into linear RNA with head-to-tail structures. **b.** The algorithm of Cirit: (1) define a 10bp fragment at the 3’ end of the transcript as the starter seed. (2) search the first seed repeat from the 5’ end of the transcript using consensus sequence alignment. (3) extract the sequence before the first seed repeat and assign it as the successor seed. (4) compare the successor seed against the fragment of same size just before the starter seed at the 3’ end. If the two fragments match well, then the transcript sequence is the potential circular RNA. (5) Remove the successor seed from the 3’ end of the transcript to yield the full-length sequence of circRNA. The BSJ sites can be determined by mapping the sequence against the reference genome, if available.

To prove the method applicable to other datasets, we also screened circRNAs in 90 public circRNA-enriched (poly(A)-/ribo-) RNA-seq datasets using Cirit. These transcriptomes were determined in 21 different cell lines of 15 distinct tissues (Additional file 4: Table S3). From these transcriptomes, we identified overall 40,747 circRNAs, of which 16,643 circRNAs had full-length sequence support. After data integration, we obtained 28,813 nonredundant circRNAs. As summarized in the Additional file 4: Table S3, the majority (80 out of 90) of transcriptomes yielded circRNAs at a low detection rate of <1% (including partially assembled circRNAs, same to all following statistics), even they were prepared in a mode of circRNA-enrichment (poly(A)^-^/ribo^-^). The circRNA detection rate had no significant propensity on cell types or tissues. However, the rate was substantially affected by the sequencing quality and sequencing length. Many (69 out of 80) transcriptomes determined in the sequencing read length of <100bp produced less than 500 sequence-supported circRNAs (the detection rates <1%). An exception was the breast cancer cell line MCF-7. Seven out of twelve MCR-7 transcriptomes determined in 51bp read length yielded 2,065 circRNAs in average, and the circRNA detection rate was about 8.49%.

In addition, we also applied Cirit to detect circRNAs from four randomly selected transcriptomes (ribo^-^) of *Arabidopsis* thaliana, *Pteropus* alecto, *Malus* domestica, and *Sus scrofa* domesticus from the NCBI. These organisms have either no genome or poorly annotated genomes up-to-date. From these transcriptomes, we identified 643, 548, 436, and 758 sequence-supported circRNAs, respectively (Additional file 4: Table S3). This result proved the Cirit method was generally applicable.

#### Experimental evaluation

According to the conventional practice, we selected 26 typical circRNAs for experimental validation in HepG2 cell line (Additional file 2: Table S1). These circRNAs covered different circRNA types. They were further categorized into three groups. Group1 included fourteen circRNAs which were identified in both HepG2 and other cell lines. Group2 included ten circRNAs which only identified in HepG2 cell line of this study once. Group3 included two circRNAs (circAFG3L1P and circNBPF14) which were not detected in HepG2 cell line but other cell lines. All PCR experiments were undertaken on the same RNA library of HepG2 cell line. As the results (Figure 2a & Additional file 1: Figure S1), 50.00% and 20.00% circRNAs were PCR-validated for Group1 and Group2, respectively, by specially-designed divergent primers, whatever the samples were treated with RNase R or not. The RNase R resistance proved the amplified RNA fragments belonging to the circRNAs. Surprisingly, circNBPF14 was also detected in the HepG2 cell line that it was supposed not to. Additional Sanger sequencing of selected PCR products further confirmed the back-splice junctions on the circRNAs (Figure 2b & Additional Information). For the 14 failed validations, 11 failed PCR amplifications whatever they were treated with RNase R or not, and three produced the wrong bands (Additional file 2: Table S1). The different validation rates for three groups suggested the PCR-validation rate was somehow related to the expression consistency, which we will discuss later.

**Figure 2.**
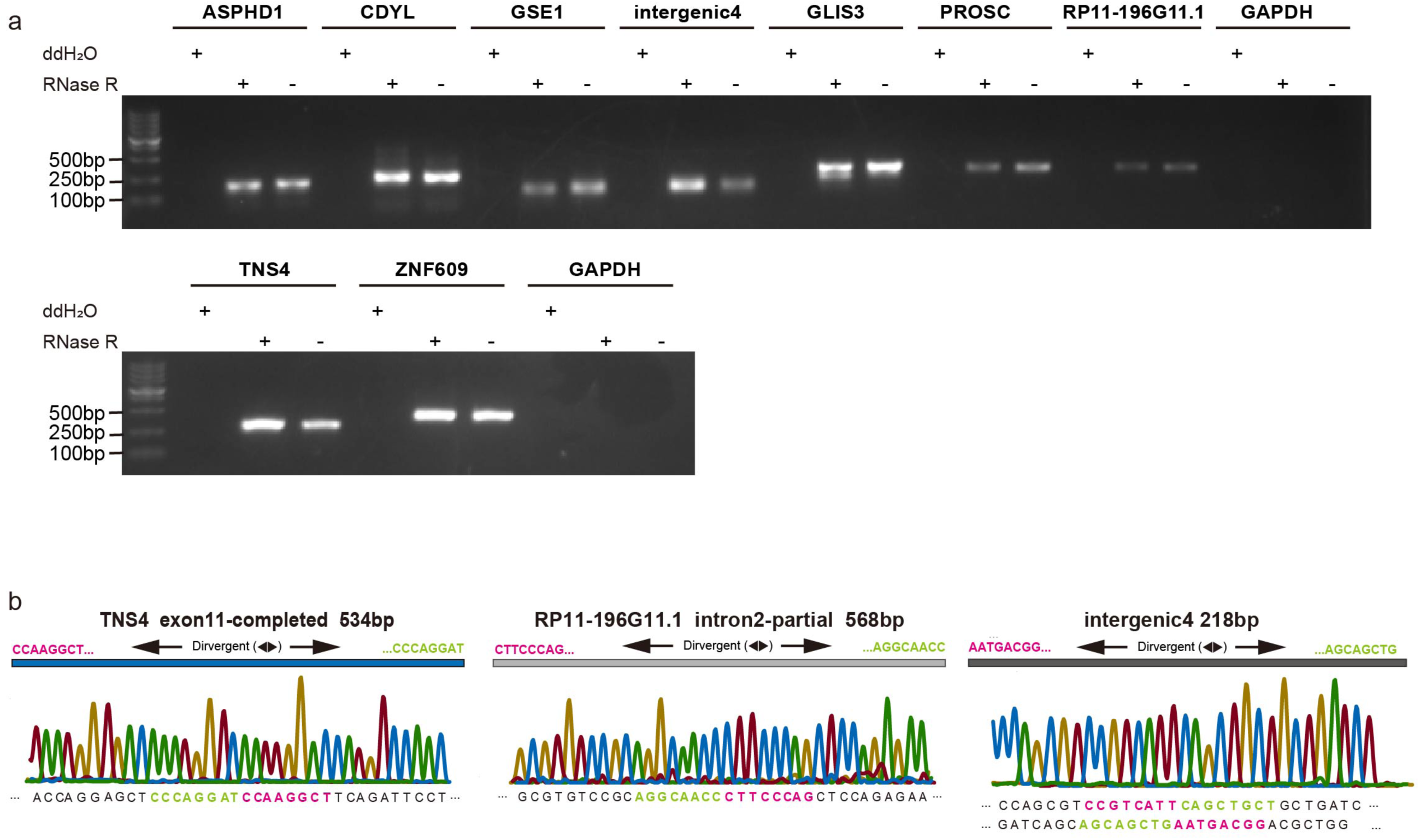
Experimental validation of selected circular RNAs. **a.** Experimental validation of selected circRNAs detected by Cirit. Each circRNA was amplified by divergent primers (crossover the back-splice junction span) in three different conditions: negative control ddH2O, sample treated with RNase R, or not. The PCR products were furtherly confirmed by Sanger sequencing. Only twelve selected experimental results were showed in this figure. **b.** Sanger sequencing of PCR products for selected circRNAs (circTNS4, circRP11, and an intergenic circRNA).

#### Performance comparison to other circRNA detection algorithms

To fairly evaluate the performance of Cirit, we compared it to five mainstream circRNA detection tools, including the annotation dependent (annotated-exon based) algorithm CIRCexplorer2 [25], the annotation independent (GT-AG based) algorithms find_circ [4] CIRI [13], and CIRC-full [24], and the combinational algorithm circRNA_finder [26] (Table 1). The comparison was made using the circRNA-enriched (poly(A)^-^/ribo^-^) transcriptome as the example dataset and the comparison results were summarized in Table 1. Cirit, CIRCexplorer2, circRNA_finder, CIRI, find_circ, and CIRC-full detected 8,182, 14,723, 23,491, 13,756, 16,233, and 13,869 circRNAs from the transcriptome, respectively. The cross-validation rate (by at least one other method) were 70.33%, 77.78%, 53.96%, 73.73%, 61.06%, and 54.67%, respectively (Table 1). The cross-validated rate by all six algorithms was only about 36.44%. These results manifested the dilemma that different methods were inconsistent in circRNA detection. The high false positive/negative rate would be a significant problem.

**Table 1.**
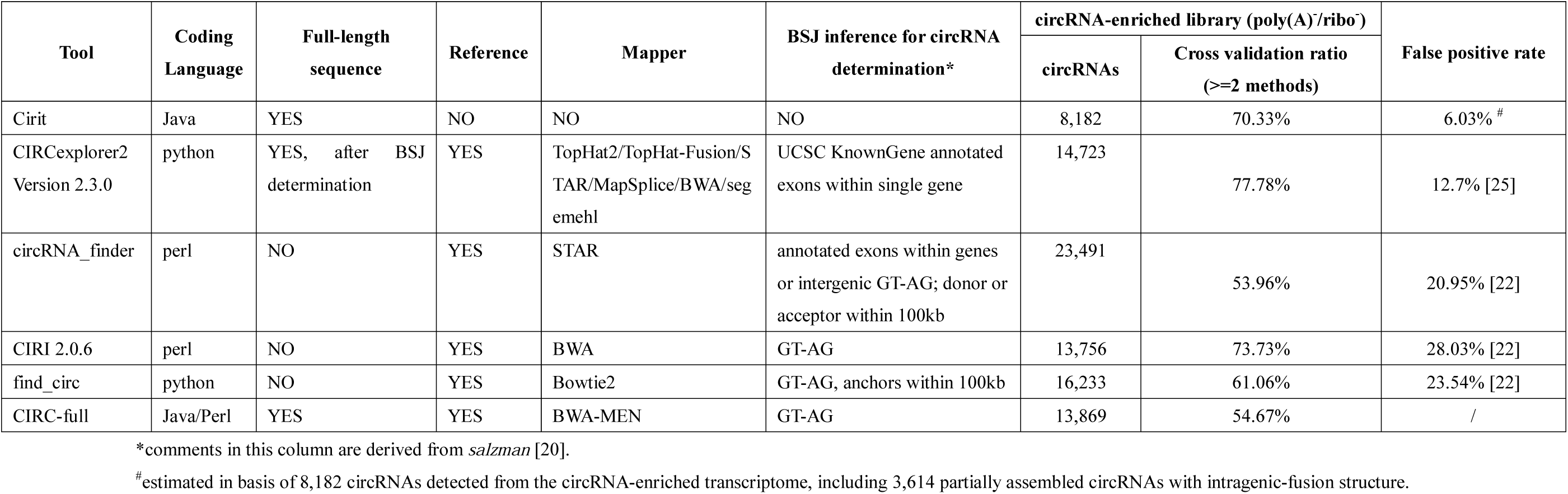
Comparison of Cirit with four selected circRNA detection methods.

In 12 experiment-validated circRNAs of this study, CIRCexplorer2 missed the intronic circRP11-196G11.1 and an intergenic circRNA (Additional file 3: Table S2). Unsurprisingly, the GT-AG based methods (find_circ, CIRI, and CIRC-full) did not detect the intronic circDUS3L since it was spliced at the non-canonical sites (Additional file 3: Table S2 & Additional file 1: Figure S2c). According to our estimation in this study, about 45.27% of total sequence-supported circRNAs were back-spliced at the canonical patterns like CT-AG/CT-GC (we will discuss in latter section). The GT-AG based methods failed to detect them all. Surprisingly, the GT-AG based method CIRI also missed circRP11-196G11.1 and the intergenic circRNA which spliced at the typical GT-AG sites (Additional file 1: Figure S2a&b).

#### Estimation of the false positive rate and the false negative rate

Prior studies have estimated the false positive rates of circRNA detection for CIRCexplorer2 (12.7%) [25], circRNA_finder (20.95%) [22], CIRI (28.03%) [22], and find_circ (23.54%) [22]. Here, we also made a rough estimation of false positive rate for Cirit in basis of the two parallel HepG2 transcriptomes determined in this study. Since the circRNAs detected by Cirit had sequence support, the major false positives may consist of the linear intragenic chimeric transcripts and the wrongly assembled transcripts.

The linear intragenic chimeric transcripts are a kind of non-collinear transcripts (NCLs). Current circRNA detection methods are incapable of distinguishing linear NCLs from circular NCLs (i.e., circRNAs) [15, 27] since they may share the same back-splicing junction [28]. In this study, the false positive rate caused by the linear intragenic chimeric transcripts was estimated as following: We identified 4,164 NCLs from the circRNA-enriched (poly(A)^-^/ribo^-^) transcriptome using the tool NCLscan [15]. These NCLs were all circRNAs theoretically since the linear transcripts have been digested by RNase R during the library preparation. In the same way, we identified 1,711 NCLs in the parallel total RNA transcriptome, which may consist of both linear NCLs and circRNAs. Of them, 686 were mutual to both transcriptomes, they were potential circRNAs [15]. Worthy of mention, three experiment-validated circRNAs (CDYL, GSE1, and ZNF609) were in the mutual list. Excluding the 686 circRNAs, the remaining 1,025 NCLs were thus likely linear chimeras. Comparing these 1,025 linear NCLs to the 1,660 circRNAs predicted by Cirit, the overlapping 210 NCLs were considered potential linear intragenic chimeras. Therefore, the false positive rate of Cirit caused by the linear intragenic chimeras was estimated to be about 12.65% (210/1660) for Cirit when it was applied to the total RNA transcriptome. In the same way, we ferreted out 349, 346, 364, 342, and 357 linear intragenic chimeras from the circRNAs predicted by CIRCexplorer2, circRNA_finder, CIRI, find_circ, and CIRC-full, respectively. In addition, we also compared the circRNAs predicted by Cirit and other five methods against the human NCL (chimeric) transcript database ChiTaRS [29], which found 0, 34, 40, 31, 33, and 37 intragenic NCL transcripts, respectively. These results suggested Cirit outperformed other five methods in distinguishing linear chimeric NCLs.

Of 8,182 circRNAs predicted in the circRNA-enriched transcriptome, 493 sequences cannot be mapped back to the human genome with high confidence (sequence identify >95%). These circRNAs could be wrongly assembled transcripts; however, their integrity was computationally validated by read coverage. Therefore, the false positive rate caused by the wrongly assembly was estimated to be maximum up to <6.03%. Likewise, the false positive rate caused by the wrongly assembly was maximally 4.22% in the case of circRNA detection from the total RNA transcriptome. Putting all these estimation results together, the false positive rates for Cirit were <6.03% and <16.87% when detecting circRNAs from the circRNA-enriched transcriptome and the total RNA transcriptome, respectively (Table 1).

Furthermore, we also made a rough estimation of false negative rate for Cirit. The estimation assumed the circRNAs predicted from the circRNA-enriched transcriptome by other five methods but Cirit as the false negatives, which was 261 circRNAs. Since the average false positive rates for the four methods was 21.31%, about 55 out of 261 circRNAs were likely wrongly predicted (the false positives). Accordingly, the false negative rate for Cirit was roughly estimated to be about 2.46% (206/8388) if the false positives were not excluded.

#### Quantification of circRNA expression

Having the full-length sequence enabled us to use the conventional FPKM (Fragments Per Kilobase per Million mapped fragments) method to quantify circRNA expression level, instead of counting fusion reads adopted by most current methods. Here, we depicted 28,813 nonredundant human circRNA expression profiles in 91 circRNA-enriched transcriptomes using the Stringtie (v2.1.0) software. Generally, the circRNA expression levels ranged from 0.01 to 161.59, majority of which were between 2.43 to 10.25 (Additional file 1: Figure S5).

In addition, we carried out a comparison analysis on circRNA expression quantification determined by either the Stringtie or the CIRC-full. The CIRC-full is oven-fresh improved method that uses both the BSJ reads and the reverse-overlap merged reads for circRNA quantification. The comparison was made on the 1,221 mutual circRNAs detected from the circRNA-enriched transcriptome of this study by Cirit and other five algorithms. The Stringtie quantified circRNAs’ expressions in basis of circRNA sequences ranging from 3.01 to 54.67 in FPKM, the majority of which were between 5.63 to 6.27. Comparatively, the expression levels determined by the CIRC-full ranged from 0.002 to 69.75, the majority of which were between 0.57 to 1.57 in FPKM (Additional file 1: Figure S5). The comparison manifested that the Stringtie can capture higher expression signals than the CIRC-full, which may be more sensitive in quantification of circRNA expression profiles.

### Characterization of human circRNAs

#### Sequential characteristics of human circRNAs

Having sequence information enable us to re-characterize human circRNAs accurately. In overall 28,813 non-redundant circRNAs detected by Cirit, 38.98% were exonic circRNAs that consisted of only complete and partial exon sequences (Figure 3c). About 7.62% were exon-intronic circRNAs that consisted of both exon and intron elements. About 23.97% were intronic circRNAs that consisted of only intron elements. About 5.22% were intergenic circRNAs that contained no exon or intron element but intergenic fragments. The remaining 24.18% (6,966 circRNAs) were unannotated circRNAs, which cannot be mapped back to human genome with high confidence for some unknown reason (Additional file 1: Figure S3a).

**Figure 3.**
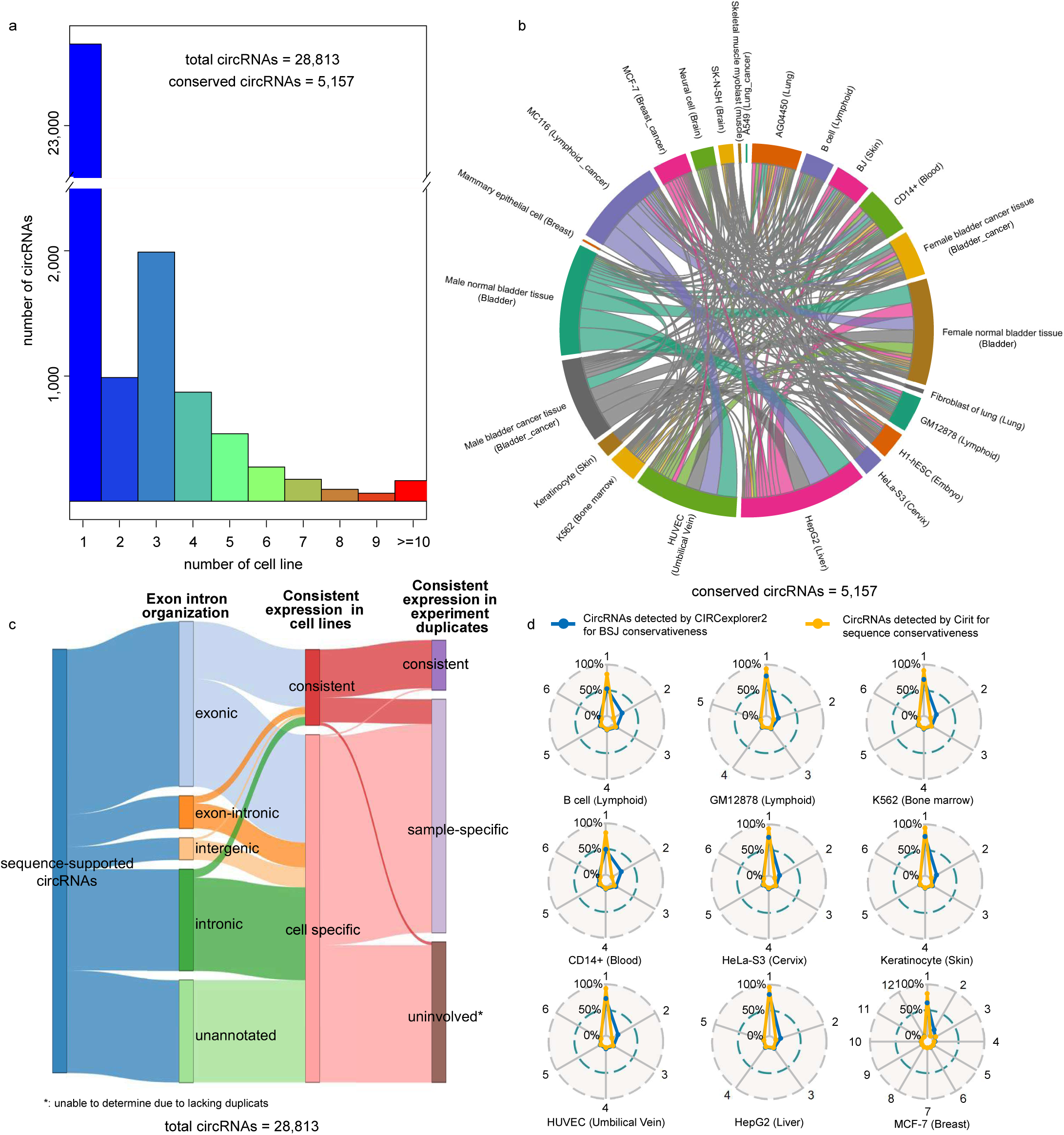
Consistency of human circRNAs over cell lines and sample duplicates. **a.** CircRNA consistency over cell lines. Only 5,157 circRNAs consistently express in at least two different cell lines with exact same full-length sequence. **b.** The majority (85.12%) of consistent circRNAs express in 2∼5 cell lines. **c.** Statistics of circRNAs by type, consistent expression over cell lines, and consistent expression in experiment duplicates. Majority of consistent circRNAs are exon-containing circRNAs. Out of 5,157 consistent circRNAs, 1,679 are also sample-specific that inconsistently express in experiment duplicates. **d.** The consistent expression of circRNAs in experiment duplicates. The analysis was undertaken in both aspects of BSJ assignment (using CIRCexplorer2) and sequence (using Cirit). Only a small portion of circRNAs are BSJ-consistent or sequence-consistent in experiment duplicates.

Almost all annotated circRNAs possessed good collinear exon-intron structures against human genome. We observed few reverse/inverse exon-intron combination, exon duplication, or extensive exon. There were also few complete exon deletions. However, we did observe partial exon deletion or partial/complete intron deletion. These findings suggested that most circRNAs likely still followed the same exquisite splice procedure in producing linear protein-coding RNAs.

The exon-containing circRNAs (both exonic and exon-intronic types) were usually composed of 1-6 exons, averagely 2.7 exons per circRNA (Additional file 1: Figure S3b). The majority (about 85%) of the circRNA exons kept intact structure. We did not observe significant preference of exon usage between circRNAs and linear protein-coding mRNAs, whatever in exon size or GC content (Additional file 1: Figure S3c). Besides, only about 25% introns were retained with their counterpart exons. We analyzed the retained introns and their counterpart exons; no significant sequence pattern was identified (Additional file 1: Figure S4a).

The length of circRNAs ranged from 200 nucleotides (nt) to 600nt in average (Additional file 1: Figure S3d), which is much shorter than that of long non-coding RNAs (lncRNA) (about 1,000nt in average [30]). Only 2,386 out of 28,813 (about 8.28%) circRNAs were longer than 1,000nt. There was no significant length difference between five types of circRNAs. In this study, the longest circRNA was an unannotated circRNAs of 92,632nt in length, its validity desired further experimental evidence. The shortest one was an exonic circRNA of 156nt in length, annotated as circBRIX1.

The GC content of circRNAs ranged from 30% to 70%, averagely 46.22% (Additional file 1: Figure S3e). Different circRNA types were slightly different in GC content. In general, the exon-containing circRNAs had comparatively lower GC content than that of the intronic, intergenic, and unannotated circRNA. There was no significant GC-content difference between circRNAs and lncRNAs; however, the circRNAs had significantly lower GC content than that of protein-coding RNAs in average (Additional file 1: Figure S3f).

#### Alternative splicing pattern and circRNA isoforms

To explore the alternative splicing pattern of circRNA generation, we mapped the nonredundant 28,813 circRNA sequences back to the human reference genome (GENECODE, GRCh37.p13). Sequence motif analysis on both BSJ flanks illustrated that circRNAs were comparatively flexible in alternative splicing than protein-coding genes (Additional file 1: Figure S4d). Other than the canonical splicing pattern of GT-AG, about 54.73% of circRNAs also used non-canonical splicing patterns such as GC-AG (0.56%) and CT-GC (0.55%). Furthermore, we made a statistical analysis on splicing motif by circRNA types. Out of 13,045 circRNAs that used canonical GT-AG splicing pattern for back-splice, 67.91% were exonic circRNAs, 11.11% were exon-intronic circRNAs, 16.61% were intronic circRNAs, and 4.37% were intergenic circRNAs. For the remaining 15,768 circRNAs that used non-canonical splicing pattern, the composition of exonic, exon-intronic, intronic, and intergenic types were 73.98%, 12.11%, 10.70%, and 3.21%, respectively.

The circRNA isoforms were defined as circRNAs originated from the same parental genes (same chromosomal loci) in regardless of BSJ selection. In this study, we identified total 17,031 circRNA isoforms for 4,387 parental genes (about 3.9 isoforms per gene) (Additional file 1: Figure S4b). We compared the circRNA isoforms of this study with the circRNA database CIRCpedia, a comprehensive circRNA database powered by CIRCexplorer2 [25]. The comparison was made to 90 transcriptome datasets (covering nine cell lines) that were studied by both this study and the CIRCpedia. From these transcriptomes, Cirit predicted overall 3,628 circRNAs isoforms from 1,353 parental genes (about 2.68 isoforms per gene in average); comparatively, the CIRCpedia database identified 59,137 circRNA isoforms for 7,241 parental genes (about 8.17 isoforms per gene in average). In CIRCpedia, 547 genes even have more than 20 circRNA isoforms (Additional file 1: Figure S4c). Between CIRCexplorer2 and this study, there were 1,305 mutual circRNA isoforms (corresponding to 853 parental genes), of which only 35.25% (460 circRNAs) were consistently detected in multiple (>2) cell lines and 25.82% (337 circRNAs) were found in single transcriptome.

### Expression consistency of circRNAs

#### Consistent expression over cell lines/tissues

Of 28,813 nonredundant circRNAs (with exact same sequence), only about 17.90% consistently expressed in at least two different cell lines, they were consistent circRNAs in sequence (Figure 3b). The remaining 82.10% of circRNAs were only detected in single cell line (Figure 3a), which were cell-specific. Among these 5,157 consistent circRNAs, about 85.05% (4,386 circRNAs) expressed in 2-5 cell lines, about 11.83% (610 circRNAs) expressed in 6-9 cell lines, and 3.12% (161 circRNAs) were highly consistent in more than ten difference cell lines (Figure 3a). In particular, circCDYL that originated from the parental gene Chromodomain Y like (CDYL) exon3 (chr6: 4891947-4892613) expressed in almost all cell lines except the endothelial cell of umbilical vein, H1-hESC, lung fibroblast cell, mammary epithelial cell, and skeletal muscle myoblast. This circRNA was also experimentally validated in mouse in previous studies [7, 31].

The 5,157 consistent circRNAs consisted of 3,902 (75.66%) exonic circRNAs, 500 (9.70%) exon-intronic circRNAs, 583 (11.31%) intronic circRNAs, and 172 (3.33%) intergenic circRNAs (Figure 3c). The composition nearly agreed with the high experiment-validation ratio (about 80%) of exonic circRNAs (Additional file 2: Table S1). Moreover, 13.61% of 5,157 consistent circRNAs used non-canonical splicing sites for back-splice. To the contrast, 46.6% of 15,874 cell-specific circRNAs used non-canonical splicing sites. These findings implied that circRNAs might adopt multiple splice machinery, in which the consistent circRNAs may prefer to using canonical alternative splice mechanism for robust genesis.

#### Consistent expression in experiment duplicates

The poor expression consistency of circRNAs was observed not only between different cell lines but also between duplicated samples. We selected nine distinct cell experiments (covering eight tissues) that had at least five experiment duplicates for circRNA expression consistency analysis, using CIRCexplorer2 and Cirit for BSJ assignment and sequence reconstruction, respectively. As the results, only a small portion (maximally 5.17%) of circRNAs were BSJ-consistent in all duplicated samples; the ratio of sequence-consistent circRNAs were mostly under 2% (Figure 3d & Table S4). To the contrast, majority of circRNAs were just sample-specific (Additional file 5: Table S4). Interestingly, we found that about 32.5% (1,679 out of 5,157) of the consistent circRNAs were also sample-specific (Figure 3c). It was likely that the low expression causes the inconsistent capture of circRNAs by current experiment procedures. In twelve experiment-validated circRNAs of this study, nine were consistent circRNAs. For the fourteen experiment-failed circRNAs, eleven were sample-specific circRNAs or not involved in consistency analysis due to lacking of duplicates (Additional file 2: Table S1). These results manifested that most of circRNAs were likely produced spatiotemporally, and the production of circRNAs may not follow a strict mechanism.

## Discussions

### *De novo* sequence-based circRNA detection provides a straightforward and universal way for confident circRNA research

Current detection of circRNA is grounded on seeking sequences or reads crossover a pair of reverse splicing sites [19]. Such strategy highly depends on good genome reference and accurate exon annotation. In practice, the imperfectness of genome annotation and the complexity of alternative splicing make high false positive rate a problematic issue in circRNA detection [20]. Although, the GT-AG based methods show some advantages in annotation-free circRNA detection, this study as well as several previous studies [4, 32-35] found prevalent non-canonical splicing events in circRNA generation. Another setback is that nowadays circRNA research barely present full-length sequence in advance. Without sequence information, many conclusions made for circRNAs, including the BSJ assignment, sequential characteristics, biogenesis, and biological functions, could be incomplete, unilateral, and even misleading. Furthermore, ascribing to the blind spots of BSJ-based algorithms themselves [20] and current experimental boundedness, accurate evaluation of false positive/negative rate in genome-wide circRNA detection is still an unsolved problem. Hence, there is a huge demand of new algorithms that can jump out of the conventional frameworks constrained by current circRNA detection methodologies. The Cirit method introduced in this study is just the one.

To our limited knowledge, Cirit is up-to-date the first reference-free, *de novo* circRNA detection algorithm that provides direct sequence evidence for RNA circularization. It breaks the dependency of good quality genome in circRNA detection, and thus allows easy deployment to any organism, even under the circumstance of draft genome or no genome. The reconstructed sequence to large extent avoids the complexity and uncertainty in BSJ assignment. However, some circRNAs still cannot be consistently validated by experiments as what we met in this study. The failure could attribute to several factors like circRNA expression consistency, RNA stability, low expression, RNA degradation, primer design, and other experiment procedures. In particular, the poor circRNA expression consistency between cells and sample duplicates could be one of the major factors.

Of course, the Cirit has its weakness. It relies on sequencing quality and subsequent *de novo* assembly, which weakens its power in detecting the full profile of circRNAs in a sample. The Cirit could miss a fraction of circRNAs due to the limitation of *de novo* assembly in reconstructing sequence with head-to-tail overlapping structure. However, mapping transcript sequence back to the reference genome, if available, can partially recover these missed circRNAs by identifying the intragenic-fusion structure. After all, the estimated false negative rate is only about 2.46%. Besides, the Cirit outperforms current methods in distinguishing chimeric NCL transcripts from circRNAs. Regretfully, it still cannot fully eliminate the false positives caused by the repeat-contained transcripts like array chimeras. The weakness attributes to the flaw of *de novo* assembly itself. Although, current version of Cirit is yet powerful enough to serve as a benchmark for genome-wide circRNA detection, its reference-free peculiarity makes it a good control for probe design and further functional exploration of circRNAs.

### Neat exon-intron organization and poor expression consistency of circRNAs challenge their role in precise posttranscriptional regulation

Early studies have identified a number of circRNAs in different organisms and subsequently concluded several sequential features of circRNAs[4, 36]. However, these analyses grounded on the circRNA sequences inferred by BSJ assignment and genome annotation, which may not be always accurately interpreted. In this study, we re-characterized human circRNAs according to the direct circRNA sequences.

In general, the circRNAs are 200-600nt in length. Like lncRNAs, the circRNAs have wide ranges of GC content from 30% to 70%. Most circRNAs keep good collinear exon-intron structure against human genome. There are few reverse or inverse exon/intron combination structures. The majority of human circRNAs are exonic types, containing 1-6 exons, averagely 2.7 exons per circRNA. This amount agrees with recent report of 1-5 exons in animal circRNAs [4]. Many exonic circRNAs consist of two or three exons instead of one exon. There are few exon-missing events during circularization; however, partial deletion of exon sometimes occurs. The circRNAs prefer complete exon (about 85%) rather than partial exon. Just as reported in prior work [36], there somehow exists few exon duplication or extensive exon. The intron of circRNA is sometimes, only about 25% in this study, retained with the counterpart exon. Comparing to the linear protein-coding mRNAs, the circRNAs do not have significant preference of exon usage such as exon size and GC content. This result agrees with prior finding [37]. The neat exon-intron organization hints circRNAs are likely spliced via an exquisite machinery.

Contrast to the fine sequential organization, circRNAs are mostly inconsistent in expression. For the first time, we unveil the expression consistency of circRNAs over cell lines and sample duplicates, from both aspects of BSJ assignment and sequence. We affirm circRNAs are heterogeneous in sequence and expression. The majority of human circRNAs are sample-specific circRNAs. Only a very small portion of circRNAs are BSJ-consistent (maximally 5.17%) and sequence-consistent (under 2%) in sample duplicates. About 17.90% of circRNAs in this study are consistently express in multiple (>2) cell lines; however, about 32.56% of these consistent circRNAs are sample-specific. This may partially answer for the low experiment-validation rate and the poor reproducibility of circRNA experiments.

Further motif analyses manifest about 45.27% of human circRNAs use the canonical GT-AG motif for back-splice, while the remaining 54.73% of circRNAs use plenty of non-canonical patterns like GC-AG/CT-GC, instead. Therein, more than 86% of consistent circRNAs and less than 50% of sample-specific circRNAs uses canonical splicing site for genesis. More than 78% of exonic circRNAs uses canonical motifs for BSJ generation; comparatively, more than 68% of intronic circRNAs cut at the non-canonical splicing motifs. Putting these data together, we speculate circRNAs might have heterogeneous mechanisms in biogenesis and regulation. CircRNAs could produce via any or combination of lariat-driven circularization [38], intron-pairing-driven circularization [11, 39, 40], RBP pairing driven circularization [38, 41], and other undiscovered mechanisms. In particular, consistent circRNAs and sample-specific circRNAs likely response to different regulatory signals, produce in different mechanisms, and execute different biological functions. Therein, consistent circRNAs may still follow the canonical transcription mechanism as linear RNAs in most cases. They express in multiple cells or tissues and are capable to execute established functions. However, sample-specific circRNAs intend to produce in an optional and flexible way, which is likely time and space dependent. Therefore, research on sample-specific circRNAs are sometimes hard to repeat. From the point of circRNA heterogeneity, we have reason to question the ability of most circRNAs in precise posttranscriptional regulation, which substantially request a consistent and robust circRNA production. Further experimental validations are desired.

## Conclusions

In conclusion, current understanding of circRNA is still limited and fragmented. Identification of transcript circulation introduced in this study provides a straightforward and universal way for reliable circRNA research. It largely boosts the successful rate in designing highly specific probes to monitor circRNA behavior in cell. In particular, the reference-free solution brings circRNA research to the organisms that have no genome or poorly annotated genome. This will substantially accelerate our functional understanding of circRNAs.

## Materials and methods

### Cell culture and RNA isolation

The human HepG2 cells were cultured in the 60mm plate medium of DMEM (#SH30045.04, Hyclone, USA) with 10% FBS (#A18156, Moybio, China) in incubator (37°C, 5% CO2). The grown cells were lysed for total RNA extraction using the TransZol Up kit (#ET111, TransGen) according to the manufacturer’s instruction. Before sending for sequencing, we checked the RNA quality using the Agilent 2200 TapeStation (Agilent Technologies, USA). Subsequently, we depleted the ribosomal RNAs (rRNAs) in the total RNA samples using the Epicentre Ribo-Zero rRNA Removal Kit (illumina, USA).

### Library preparation and sequencing

In this study, we prepared two RNA libraries in parallel from the same RNA sample: the total RNA library (ribosomal RNA depleted, ribo^-^) and the circRNA-enriched library (linear RNA and poly(A) depleted, poly(A)^-^/ribo^-^). To prepare the circRNA-enriched library, we treated 5μg total RNA (ribo^-^) with 20U RNase R (#RNR07250, Epicentre, USA) at 37 °C for 30 minutes to deplete linear RNAs. Eventually, 1μg rRNA-depleted total RNAs and 2μg RNase R treated RNAs were used for library preparation. For each of these two libraries, the RNAs were fragmented into about 300bp for cDNA generation. Size selection was performed before cDNA adaptor ligation according to the guidance of NEBNext® Ultra™ RNA Library Prep Kit for Illumina. The library products were evaluated using the Agilent 2200 TapeStation and Qubit®2.0 (Life Technologies, USA), and then diluted to 10pM. Clusters were generated on the HiSeq3000 pair-ended flow cell, and the samples were then sequenced on HiSeq 3000/4000 PE Cluster Kit with the read length equal to 150bp.

### RNA sequencing data preprocessing and *de novo* assembly

Before proceeding the transcriptome assembly, we performed quality control on the raw sequencing data to exclude low-quality reads, including adaptor-contained reads, reads with N-content >10%, and other low-quality reads that own more than 50% bases with sQ <=5. In basis of the clean sequencing data, we adopted the IDBA-Trans (version 1.1.1) [42] software for *de novo* transcriptome assembly in this study. We chose the IDBA-Trans tool because they have advantages of less computational time, more sensitivity in detecting low-expressed transcripts, and more efficiency in regenerating full-length sequence [42]. Other *de novo* methods like Velvet [43] also work well. Worthy of mention, not all *de novo* methods were suitable for circRNA assembly, especially those methods that simply removed tips or bubbles in the *de Bruijn* graph computation. For transcriptome assembly by IDBA, a series of kmer sizes (from 100 to 150, step=5) were tried to elect the right kmer, which yielded the largest transcript number and the longest N50.

The raw sequencing datasets determined in this study have been deposited in the Genome Sequence Archive [44] in BIG Data Center, Beijing Institute of Genomics (BIG), Chinese Academy of Sciences, under accession numbers CRA001233. The data are publicly accessible at http://bigd.big.ac.cn/gsa.

### Reference-free detection of circRNAs and annotation

In this study, we developed a new algorithm for reference-free circRNA detection from transcriptome. In principle, the algorithm searches the transcript containing the head-to-tail overlapping structure that allows the transcript circularized. As illustrated in Figure 1, the algorithm takes five continued steps to detect circRNAs in the *de novo* assembly transcriptome: (1) define a 10bp fragment at the 3’ end of the transcript as the starter seed; (2) search the first seed repeat from the 5’ end of the transcript by consensus sequence alignment; (3) extract the sequence before the first seed repeat and take it as the successor seed; (4) compare the successor seed against the fragment of same size just before the starter seed at the 3’ end. If the two sequences match well, then the transcript is the potential circular RNA. (5) Remove the sequence after the successor seed from the 3’ end of the transcript to yield the full-length sequence of circRNA.

We annotated the circRNA and determined the back-splice junction by mapping the sequence to the human genome (GENECODE, GRCh37.p13) with GMAPv3.0 [45]. The start and the end of back-splice sites correspond to the smallest and the largest coordinates, respectively. For convenience, we coded the circRNA detection algorithm as a standalone software, namely Cirit, using the Java language. The Cirit software has been tested, packed, and deposited at http://www.bio-add.org/CIRIT/ or http://bioinf.xmu.edu.cn/CIRIT/ for free downloading.

In addition to the internal transcriptomes, we also used Cirit to detect circRNAs from 90 external transcriptomes determined for 22 different human cell lines and four transcriptomes determined for organisms such as *Arabidopsis* thaliana, *Pteropus alecto, Malus domestica*, and *Sus scrofa domesti*cus. These transcriptomes were sequenced in total RNA modes. The raw RNA-seq datasets were downloaded from either ENDCODE or NCBI SRA (Additional file 4: Table S3).

### Circular RNA validation

We digested 5μg total RNA extracted from the same cultured human HepG2 cells using 20U of RNase R (#RNR07250, Epicentre, USA) at 37 °C for 30 minutes to deplete linear RNAs. Subsequently, the circRNAs were recovered according to the method described by *Anna* [40]. The circRNAs were reversely transcribed into cDNAs by occasional primers according to the manufacturer’s instruction (#AT311, TransGen, China). Taking the cDNAs as templates, we performed the PCR amplification reactions by the specially designed divergent primers listed in Additional file 6: Table S5. The divergent primers were designed to produce sequence that can bridge across the BSJ site. The PCR products were checked for band size on agarose gel. Furthermore, the PCR products that satisfying the expected band sizes were validated using the Sanger sequencing (Additional information).

## Additional files

**Additional file 1:**

**Figure S1.** Experimental validation of selected circular RNAs.

**Figure S2.** The genomic information of three selected experiment-validated circRNAs.

**Figure S3.** The sequential characteristics of human circRNAs.

**Figure S4.** Alternative splicing events in circRNAs.

**Figure S5.** The expression level of sequence-based circRNA.

**Additional file 2:**

**Table S1.** The experimental validation of 26 selected circRNAs. (DOCX 24kb)

**Additional file 3:**

**Table S2.** The experiment-validated circRNAs predicted by different methods. (DOCX 19kb)

**Additional file 4:**

**Table S3.** The accession number of ENCODE RNA-seq data analyzed during the current study. (XLSX 20kb)

**Additional file 5:**

**Table S4.** CircRNA expression consistency in duplicated samples. (DOCX 16kb)

**Additional file 6:**

**Table S5.** Divergent primers for the experimental validation of circRNAs in this study. (DOCX 22kb)

## Acknowledgements

Financial support from the National Key Research & Developmental Program of China (2018YFC1003601) and the Natural Science Foundation of China (NSFC# 31671362) are gratefully acknowledged. We also appreciate valuable comments from Professor Zefeng Wang.

## Author contributions

Y.Q. and T.X. wrote the manuscript text, demonstrated majority of the bioinformatics analysis and prepared most of the figures and tables. Q.H. annotated circular RNA and packaged the algorithm into java. K.L. did the statistical analysis of circRNA length and GC content. J.D., L.C., X.Y. and F.D. prepared some of pictures. W.L., Q.J., M.L., M.C., and T.T. supplied HepG2 RNA for sequencing and did the circular RNA experimental verification. T.T. designed and presided the experimental parts of this study, reviewed the manuscript. Z.J. presided over the whole project, designed the experiments, reviewed and revised the manuscript. All co-authors have reviewed the manuscript.

## Availability of data and materials

All data used in this study are publicly available. The accession number of ENCODE RNA-seq data analyzed during the current study can be obtained from Additional file 4.

