## Additional file 1 for "Reference-free and *de novo* Identification of Circular RNAs"

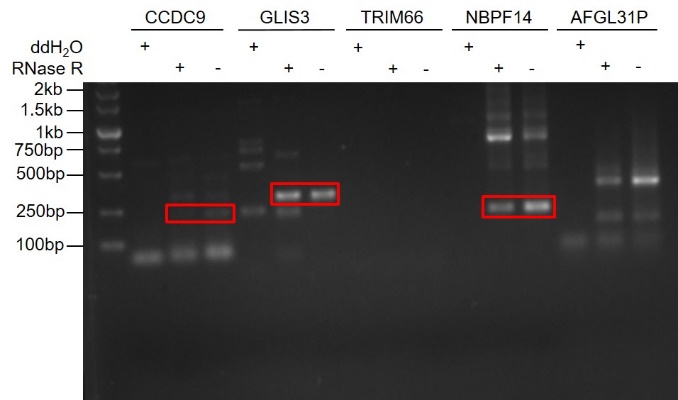

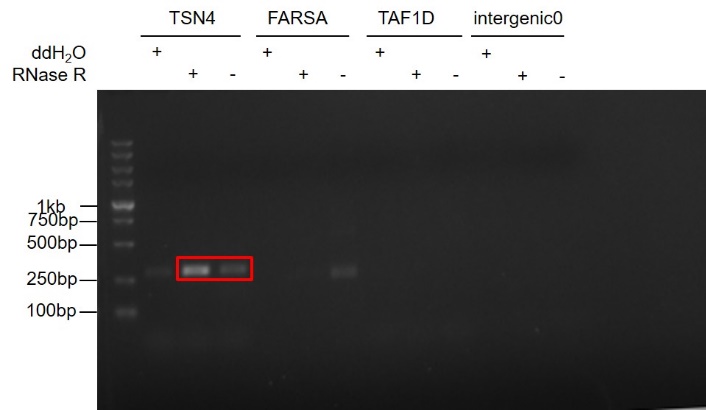

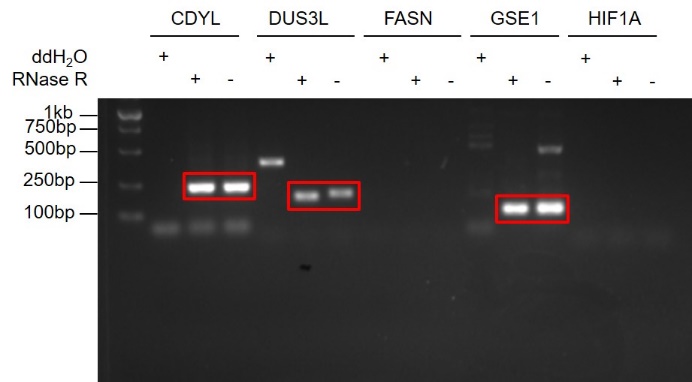

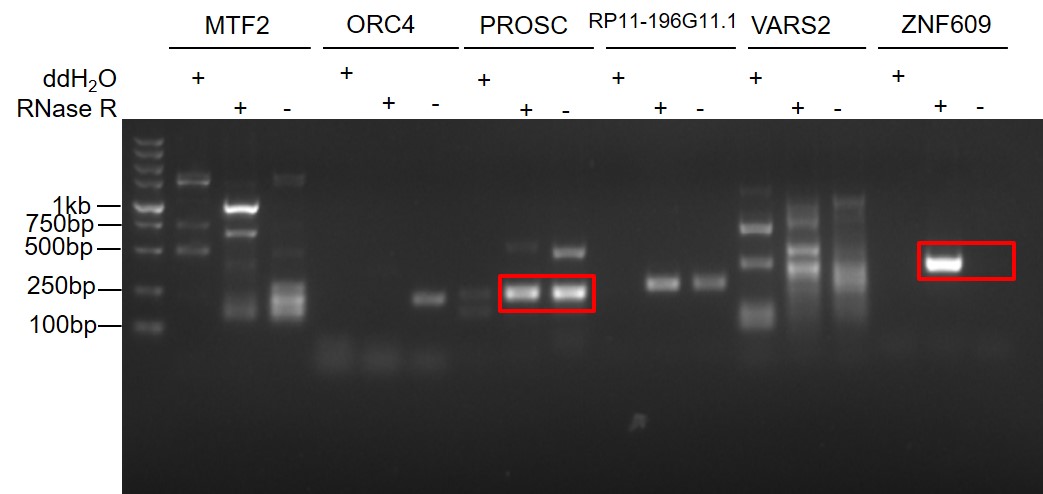

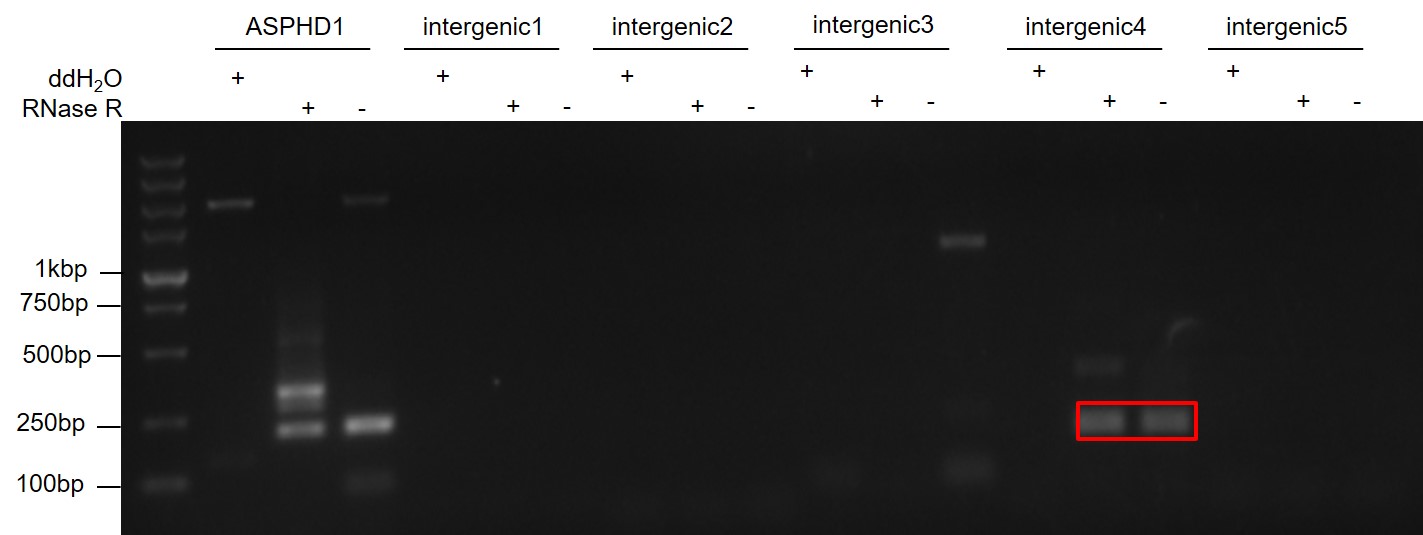

**Figure S1. Experimental validation of selected circular RNAs.** The divergent primers were designed specially to crossover the back-splice junction spans on circRNAs under three conditions: negative control ddH_2_O, sample treated with RNase R, or not. The red box stands for the expected PCR products. Sanger sequencing of the selected PCR products were given in the Additional file 6.

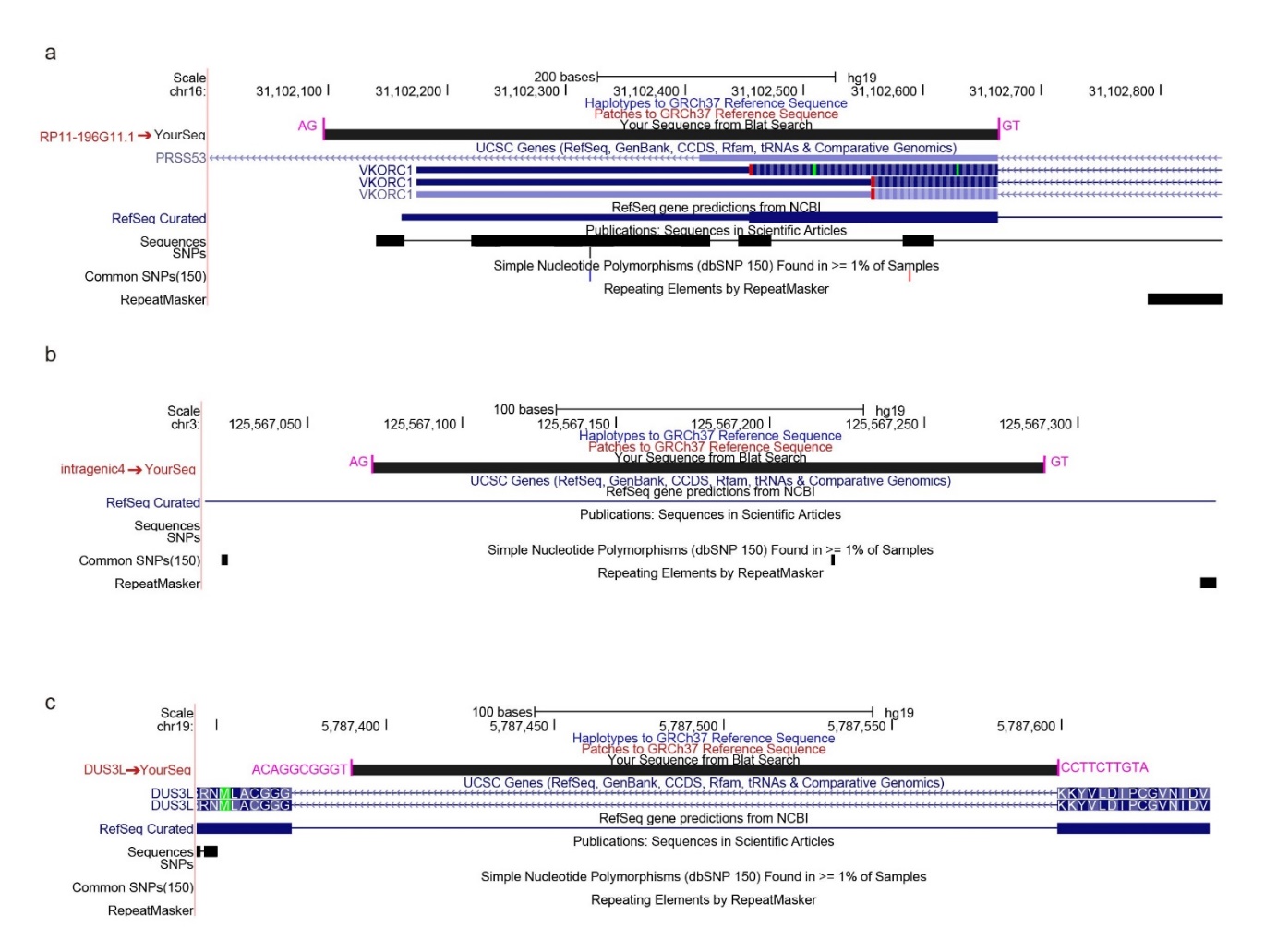

**Figure S2. The genomic information of three selected experiment-validated circRNAs**. The genome mapping used the UCSC hg19 as reference. (a) The genomic information of intronic circRNA circRP11-196G11.1. (b) The genomic information of an intergenic circRNA. (c) The genomic information of intronic circRNA circDUS3L.

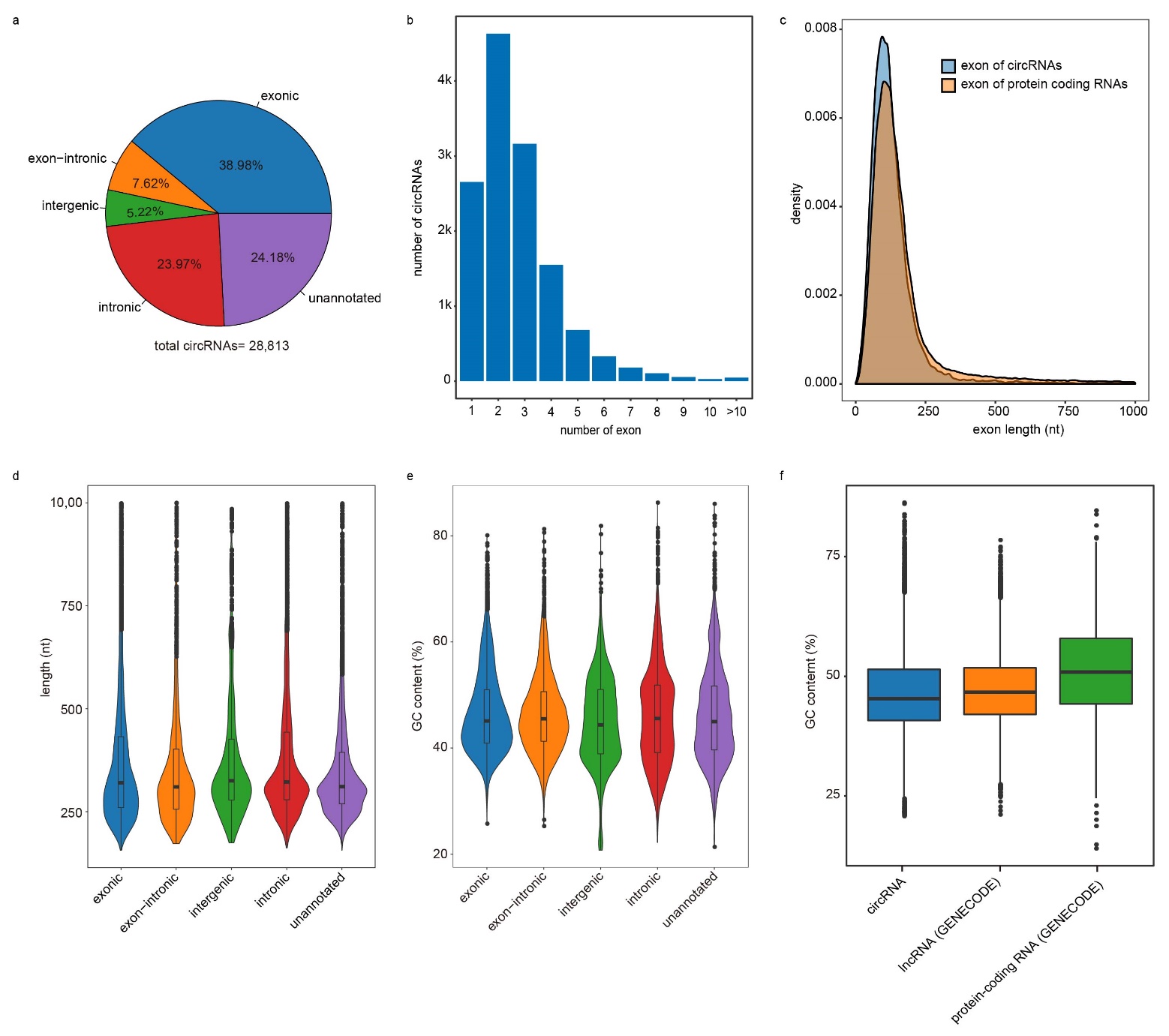

**Figure S3.** **The sequential characteristics of human circRNAs.** (a)The circRNAs detected by Cirit from 91 transcriptomes were classified into five types according to annotation: exonic, exon-intronic, intronic, intergenic, and unannotated. (b) Statistics of exon number in the exon-containing circRNAs. Majority of the exon-containing circRNAs were composed of 1-6 exons. (c) The comparison of exon length in circRNAs and protein-coding RNAs. (d) The circRNA length. Most of circRNAs ranged from 200nt to 600nt in length. (e) GC content of circRNAs ranged from 30% to 70%. Different circRNA types were slightly different in GC content. (f) GC content comparison of circRNAs, lncRNAs, and protein-coding RNAs. The protein-coding RNAs had comparatively higher GC content than that of noncoding RNAs (circRNAs and lncRNAs).

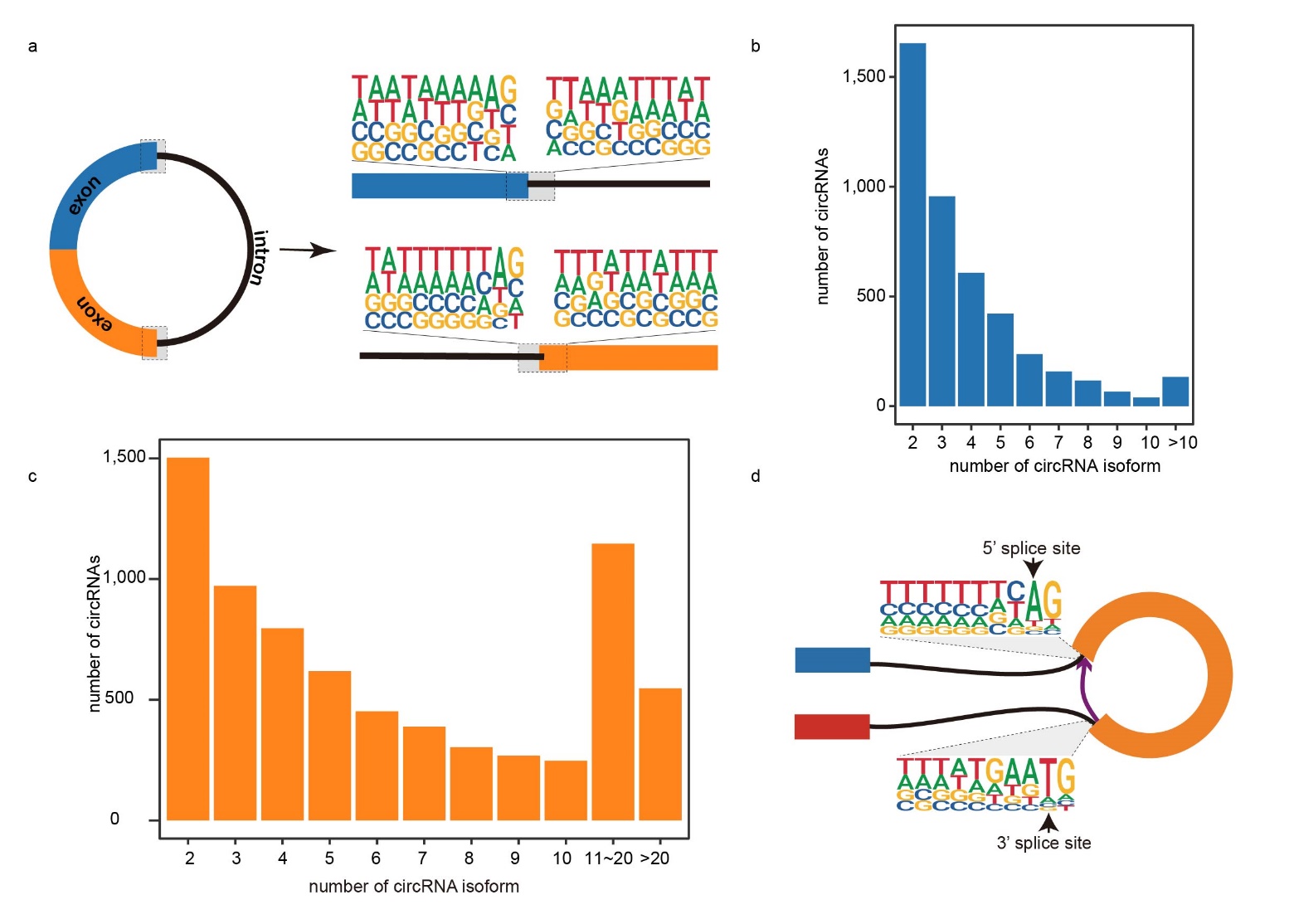

**Figure S4. Alternative splicing events in circRNAs.** (a) Sequence analysis of intron-retained circRNAs. No significant sequence pattern was identified on the boundary between exon and intron. (b) Statistics of circRNA isoforms detected by the Cirit. Total 17,031 circRNA isoforms were detected for 4,387 parental genes. Most of parental genes possess 2~5 circRNA isoforms. (c) The statistics of circRNA isoforms collected by CIRCexplore2. (d) Motif analysis of circRNA flanking sequence. Other than the canonical splice site GT-AG, non-canonical splice site like GC-AG and CT-GC were also often used by circRNAs.

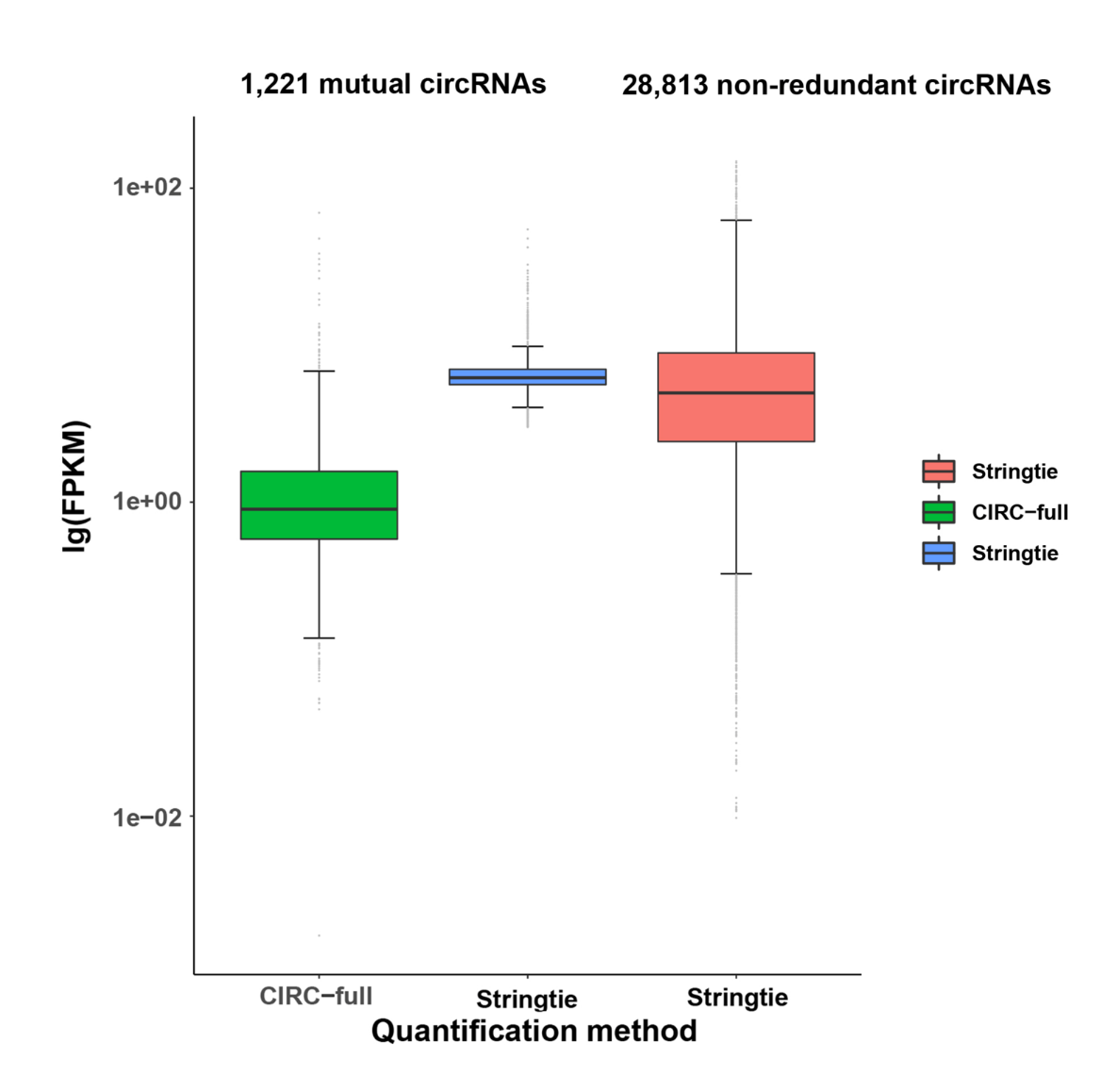

**Figure S5. The expression quantification of human circRNAs**. The analysis was made on the 1,221 mutual circRNAs detected from the circRNA-enriched transcriptome of this study by Cirit and other five algorithms. The Stringtie quantified circRNAs’ expression ranging from 3.01 to 54.67, the majority of which were between 5.63 to 6.27 in FPKM. The CIRC-full quantified circRNAs’ expression ranging from 0.002 to 69.75, the majority of which were between 0.57 to 1.57 in FPKM. The comparison manifested that the Stringtie can capture higher expression signals than the CIRC-full. The expression levels of 28,813 nonredundant human circRNAs were also determined by the Stringtie, ranging from 0.01 to 161.59, the majority of which were between 2.43 to 10.25.
