## Additional file 2 for "Reference-free and *de novo* Identification of Circular RNAs"

**Table S1. The experimental validation of 26 selected circRNAs.**

| **Group** | **CircRNA** | **Type** | **Consistent in cells** | **Sample- specificity^#^** | **CircRNA-enriched library** | **Total RNA library** | **PCR product** | **BSJ confirmed by Sanger sequencing** |
| --- | --- | --- | --- | --- | --- | --- | --- | --- |
| group3 | AFG3L1P * | exonic |  | N.A. |  |  | √ |  |
| group2 | ASPHD1 | exonic |  | N.A. | √ |  | √ | √ |
| group1 | CCDC9 | exonic | √ |  |  | √ | √ | √ |
| group1 | CDYL | exonic | √ |  | √ | √ | √ | √ |
| group2 | FARSA | exonic |  | N.A. | √ |  |  |  |
| group1 | GLIS3 | exonic | √ |  | √ | √ | √ | √ |
| group1 | GSE1 | exonic | √ | √ | √ | √ | √ | √ |
| group3 | NBPF14* | exonic |  | N.A. |  |  | √ | √ |
| group1 | ORC4 | exonic | √ |  | √ |  |  |  |
| group2 | TNS4 | exonic |  | N.A. | √ |  | √ | √ |
| group1 | ZNF609 | exonic | √ |  | √ | √ | √ | √ |
| group1 | HIF1A | exon-intronic | √ | N.A. | √ |  |  |  |
| group1 | MTF2 | exon-intronic | √ | √ | √ |  | √ |  |
| group1 | PROSC | exon-intronic | √ |  | √ |  | √ | √ |
| group2 | Intergenic0 | intergenic |  | N.A. | √ |  |  |  |
| group2 | intergenic1 | intergenic |  | N.A. | √ |  |  |  |
| group2 | intergenic2 | intergenic |  | N.A. | √ |  |  |  |
| group2 | intergenic3 | intergenic |  | N.A. | √ |  |  |  |
| group1 | intergenic4 | intergenic | √ | √ | √ |  | √ | √ |
| group2 | intergenic5 | intergenic | × | N.A. | √ |  |  |  |
| group1 | DUS3L | intronic | √ | √ | √ | √ | √ | √ |
| group2 | FASN | intronic |  | N.A. | √ | √ |  |  |
| group1 | RP11-196G11.1 | intronic | √ |  | √ | √ | √ | √ |
| group2 | TAF1D | intronic |  | N.A. | √ |  |  |  |
| group1 | TRIM66 | intronic | √ |  |  | √ |  |  |
| group1 | VARS2 | intronic | √ |  | √ | √ | √ |  |

* circNBPF14 were detected in two external ploy(A)-enriched transcriptomes (ENCODE experiment ID: ENCSR985KAT) but not in both transcriptomes prepared in this study.

^#^ Not available (N.A.). N.A. stands for no sufficient duplicate data available for conservativeness analysis.
