## Additional file 3 for "Reference-free and *de novo* Identification of Circular RNAs"

**Table S2.** **The comparison of experiment-validated circRNAs by different methods.**

| **CircRNA** | **Coordinate in genome (hg19)** | **Cirit** | **CIRCexplorer2** | **circRNA_finder** | **CIRI** | **find_circ** |
| --- | --- | --- | --- | --- | --- | --- |
| ASPHD1 | chr16:29917447-29917108 | √ | √ | √ | √ | √ |
| CCDC9 | chr19:47767860-47768203 | √ | √ | √ | √ | √ |
| CDYL | chr6:4891947-4892613 | √ | √ | √ | √ | √ |
| DUS3L | chr19:5787391-5787599 | √ | √ |  |  |  |
| GLIS3 | chr9:4286038-4286523 | √ | √ | √ | √ | √ |
| GSE1 | chr16:85667520-85667738 | √ | √ | √ | √ | √ |
| Intergenic4 | chr3:125567073-125567290 | √ |  | √ |  | √ |
| PROSC | chr8:37623044-37623873 | √ | √ | √ | √ | √ |
| RP11-196G11.1 | chr16:31102097-31102664 | √ |  | √ |  | √ |
| TNS4 | chr17:38652239-38652772 | √ | √ | √ | √ | √ |
| ZNF609 | chr15:64791492-64792365 | √ |  | √ | √ | √ |
