## Additional file 5 for "Reference-free and *de novo* Identification of Circular RNAs"

**Table S4. Consistent expression of circRNAs in experiment duplicates**. The circRNAs were detected by either CIRCexplorer2 or Cirit. Ten distinct cell experiments (total 58 transcriptomes) containing at least five duplicates were included in this analysis.

| **Cell lines** | **Duplicate Number** | **Consistency in BSJ**  **(CIRCexplorer2)** | | | **Consistency in sequence**  **(Cirit)** | | |
| --- | --- | --- | --- | --- | --- | --- | --- |
|  |  | Total circRNAs | Singleton (%) | Consistency in all datasets (%) | Annotated circRNAs | Singleton (%) | Consistency in all datasets (%) |
| B cell | 6 | 10,025 | 53.25 | 4.63 | 907 | 81.59 | 0.88 |
| CD14 | 6 | 13,331 | 49.55 | 5.17 | 1,169 | 81.86 | 1.71 |
| HUVEC | 5 | 9,349 | 81.21 | 0.76 | 796 | 94.85 | 0 |
| GM12878 | 6 | 22,766 | 72.51 | 0.95 | 1,396 | 92.05 | 0 |
| HeLa-S3 | 5 | 19,703 | 78.04 | 1.08 | 898 | 91.76 | 0.11 |
| HepG2 | 6 | 21,323 | 75.03 | 1.25 | 1090 | 91.65 | 0 |
| K562 | 6 | 34,168 | 71.39 | 0.86 | 1,304 | 88.65 | 0 |
| Keratinocyte | 6 | 10,876 | 76.80 | 0.45 | 626 | 92.97 | 0.16 |
| MCF-7 | 12 | 19,026 | 64.33 | 0.71 | 1,579 | 82.33 | 0.06 |
