## Additional file 6 for "Reference-free and *de novo* Identification of Circular RNAs"

**Table S5. Divergent primers for the experimental validation of circRNAs in this study.**

| **circRNA** | **circRNA type** | **upstream** | **downstream** |
| --- | --- | --- | --- |
| ASPHD1 | exonic | AGCGCCAGGCCCTCGACTTT | GGGTGCCAGAGGTCCACGAT |
| CCDC9 | exonic | ATGTTCTGCCTGCGCCGCTCCT | ATGAGCGCAACCAGCGGGAAGG |
| CDYL | exonic | TGGCGTCTGTTGAAGTCGTG | CAGGCCCAGGATACACCCA |
| GLIS3 | exonic | GGCTCGGATGGCAGGAAT | GCGATTCCAGGTCACCCA |
| GSE1 | exonic | GTCCTGGTCGCGGTGGAAA | GCGCCATCCTCCAGCTTTG |
| NBPF14 | exonic | AACAGCCAAGCCAACACG | GGAAGAAGACCAAGACCCAT |
| TNS4 | exonic | GCAGTGGGCTGGACATCA | CCCCAGAGGACCTTGACTC |
| ZNF609 | exonic | GAGCCTGACATTTCCAGTTTCT | TCCTTTGGGAACTAAACCG |
| PROSC | exonic-intronic | TTTGCTGACCGCCACTAG | ACATGGGCAGCGCACTTT |
| Intergenic4 | intergenic | GGAAGCCCTGGAAGGAGAT | CTTTGTCACGATGACCTCCCG |
| DUSL3 | intronic | TGACCCTCCACCGAGACAT | CCCTGGAGTCCCTGCCTTTC |
| RP11-196G11.1 | intronic | GAGGGAAGGTTCTGAGCAAT | GCATAGGTGGTGATACAAACA |
| GAPDH | negative control | AGAAGATGCGGCTGACTG | AGCCACATCGCTCAGACA |

**The Cirit-constructed sequences and Sanger-determined sequences of the twelve experimental validated circRNAs**.

For easily visualizing the back-splice junctions in the sequences of Sanger sequencing, the head and tail sequences (10bp) of circRNAs were highlighted with yellow and green, respectively.

***The Cirit-constructed sequences of the twelve experimental validated circRNAs.***

>ASPHD1|exon2-partial|221bp

TGTTTGGTTGTACTAGGAGAAGACATAAGGCCCTGCCCCAGCCTCTCCCATCAAGCCGTCCCTGAGCGCAGTGTCTCTCCAGCCTGGGTGTGTGAAGGGAGCACCTTCCTTCAAGGGTCTGGGGCGAAGACAAAGTCGAGGGCCTGGCGCTCAGCCCCTGCCACGTTGGGGTGCCAGAGGTCCACGATGAAGACCACTCGAGGCCCATCTTCGGGGGAGCC

>CDYL|exon3-complete|667bp

GTTGAAAGGATTGTTGACAAAAGGAAAAATAAAAAAGGGAAGACAGAGTATTTGGTTCGGTGGAAAGGCTATGACAGCGAAGACGACACTTGGGAGCCGGAACAGCACCTCGTGAACTGTGAGGAATACATCCACGACTTCAACAGACGCCACACGGAGAAGCAGAAGGAGAGCACATTGACCAGAACAAACAGGACCTCTCCCAACAATGCTAGGAAACAAATCTCCAGATCCACCAACAGCAACTTTTCTAAGACCTCTCCTAAGGCACTCGTGATTGGGAAAGACCACGAATCCAAAAACAGCCAGCTGTTTGCTGCCAGCCAGAAGTTCAGGAAGAACACAGCTCCATCTCTCTCCAGCCGGAAGAACATGGACCTAGCGAAGTCAGGTATCAAGATCCTCGTGCCTAAAAGCCCCGTTAAGAGCAGGACCGCAGTGGACGGCTTTCAGAGCGAGAGCCCTGAGAAACTGGACCCCGTCGAGCAGGGTCAGGAGGACACAGTGGCACCCGAAGTGGCAGCGGAAAAGCCGGTCGGAGCTTTATTGGGCCCCGGTGCCGAGAGGGCCAGGATGGGGAGCAGGCCCAGGATACACCCACTAGTGCCTCAGGTGCCCGGCCCTGTGACTGCAGCCATGGCCACAGGCTTAGCTGTTAACGGGAAAG

>CCDC9|exon5-exon6-complete|258bp

CCGCTCGTGCTCAAGGCCACAGCGCACCCGCTCGAAGTCGGGGCCCCCCCAGTTGCGGCCGTGCCGGCGGCTCTCCTCCCGGCGGTCCCGCTCAGACTCCTCCAGGGGCCCGCTGCGTCGCCGGGGGTCGTCCAGGAAGTTCCGCACTGGGTTGGGTTCAAGAACCCCTTCCCGCTGGTTGCGCTCATACTCGGCGATCTTCTCCATCTCCTCATTCATCTTCTCAATGTTCTGCCTGCGCCGCTCCTCCCACTCCTG

>DUS3L|intron6-complete|210bp

TGGAGGGAGGGCAGGACGTGGGCTGACCCAAGACCCCTCACCCCTGCTGCCCGGAGGCCCCCACCAGGAGACGCCGACCTGCAGGAAAGGCAGGGACTCCAGGGCCACACTTGAGCCCAAACCCCTGACCCTCCACCGAGACATCGCCGGCTGGGTCAGTGCCTCCCTGCCCAGATGGGGAAACCGAGGCACACGTCCAGGGCCGCTACG

>GSE1|exon1-complete|219bp

GCATGAGCCATGAGCCCAAGTCCCCTTCGCTAGGGATGCTTTCCACCGCGACCAGGACCACCGCCACCGTCAACCCCCTCACCCCCTCGCCGCTCAATGGCGCCCTGGTGCCCAGCGGCAGCCCCGCCACCAGCAGCGCGCTGTCGGCCCAGGCCGCGCCATCCTCCAGCTTTGCCGCCGCGCTGCGCAAGCTCGCCAAACAGGCGGAGGAGCCCAGAG

>intergenic|218bp

AATGACGGACGCTGGGGCTTTGATGGGCACCGGGTGAAATGGGCAGAGTGGCGCTTACCCGGGATGGCGGTGAAGCGGGACCGGGAGGTCATCGTGACAAAGGGTGGCATGAGGTACCTGGCCTTGACGCCCTCCCCGGCTAGACGGTCCAGATTGGGGGTGTCCACATCCTGATCCTAGTCCCAGCGGAAGCCCTGGAAGGAGATCAGCAGCAGCTG

>GLIS3|exon9-complete|486bp

TTTCCCAGGATTTGGAGGAAAAGGGCTTCCAAACTCCTGCTGCTTTGGCTTTAAAGTATGTGACCCTGACATGCCTCCAGCCTGGGTGACCTGGAATCGCGGCTTCCCATTGGTGAGCATTTGTCTCCTGGGGCTTAAGGCAGGCAGATGGATGCGGCTCTCAGCCACGTTGTTCTGAGGAGCCATCCCTCCTCCTGAGGGCATCTTGAGATGGAGGTTGTTAGCAAGGCTTGCCATAGTGGGACTCGATGTGCTGCCACAGGGCGAGGGGCCAGGAGTCCCGGAGTGGGCTCGGATGGCAGGAATGTGATGACCACTGACCATCCTAGGCCCCTGTGGGGTTCCCGATGTCCGGTGGAGACTCATGCTGCATGATCTTCCATTCATTCTGAAAAACCTGTGGCCAAGACGGTCAAATATCCAATGTCACTAATGACTCCTTTCAGGCAAAGTCCAATAAGTTATCCATGGTGTGGGTTATAAGCC

>NBPF14|exon5-exon6-complete|225bp

GGGGAGCTGTTGGATGAGAAAGAGCCTGAAGTCTTGCAGGAGTCACTGGATAGATGCTATTCAACTCCTTCAGGTTGTCTTGAACTGACTGACTCATGCCAGCCCTACAGAAGTGCCTTTTACATATTGGAGCAACAGCGTGTTGGCTTGGCTGTTGACATGGATGAAATTGAAAAGTACCAAGAAGTGGAAGAAGACCAAGACCCATCATGCCCCAGGCTCAGC

>PROSC|exon1-intron1-exon2-exon3|297bp

GATCTCCCAGCCATCCAGCCCCGGCTAGTGGCGGTCAGCAAAACCAAACCTGCAGACATGGTGATCGAGGCCTATGGACATGGGCAGCGCACTTTTGGCGAGAACTACGTAAGAGCCCTTTCCTGAAGCCCTTTGGAAGCATCATGATTGCCAGGCTTCTGACTTGTTCTGTTTTGACCTTTTAGGTTCAGGAACTGCTAGAAAAAGCATCAAATCCCAAAATTCTGTCTTTGTGTCCTGAGATCAAATGGCACTTCATTGGCCACCTACAGAAACAAAATGTCAACAAATTGATGG

>RP11-196G11.1|intron2-partial|568bp

CTTCCCAGCTCCAGAGAAGGCAACACCGAGGGAGGCCCAGCACCACAGTCCATGGCAGACACATGGTTCAGACTTGGCTGATTGATCTAAGAAACTTTATTGCTCAGAACCTTCCCTCCCTGGGCAATGGAAAGAGCTTTGGAGACCAGCCCATGGGGACAGAGTCAGAGGCACTGGGTGTAAAAAAGAGCGAGCGTGTGGCACATTTGGTCCATTGTCATGTGCGGGTATGGCAGGAGGAGGGGGTAATCTAGAAGCCCCACATCTAGGGCCTTCTAGGGACCCAGATATGCCCCCTTAGGCAAGGCTCACATGCCAAAGCAAAGCAGATGAGGTCAGCCTGGCTTGGGTTGAGGGCTCAGTGCCTCTTAGCCTTGCCCTGGGGTTCTTGGACCTTCCGGAAACTGAGCCACATCAGGCTCACGTTGATAGCATAGGTGGTGATACAAACAATGCAGAAATCATAGAGCACGAAGAACAGGATCCAGGCCAGGTAGACAGAACCAGCGAGAGACACCAGGGAGCTCAGCAGCATCAGGACAGAGGCCCAGCGTGTCCGCAGGCAACC

>TNS4|exon11-complete|534bp

CCAAGGCTTCAGATTCCTCCTTCTTTCTCATTGACATGGTGCTCTGGGCCAGCTCAGCCTGGGAGCCCCCAGTCCCTGGGGGAAGCAGCTGGAAGGTGGGGTCCAGTTCCAGGATCATCTGATTGAGGCTCTCCAGTGAGAAGTCAATGTAGGAGTCAAGGTCCTCTGGGGTCCCCAAGGCCTTCTCACCAGGGGACGGCAGGAAGCAGGTGGCTTTGGCCTCCACCTGTGGGGCTTGCTGGAGTCGGCCAGGGGGCCCCATGCAGGGCACGGGGGCCATCAGGGCCTGGGCTCCCCAGCCTTCCGTGGTGTAGTAAGAACACTGGGGTGGCAGGCTGGGGCTGGGTGCTGGGTGCAGGGTCCTCCTGGGCTCATCACAAGGCGCCAAGCTGACAGCATGGCCTCCTGCCAGCAGTGGGCTGGACATCACCTGGGACATGGTGGGGGTGGTGACCTCTGCAGTTTACCTCTGGTCTTCAACCAGCCTCACTGACATCCCAGAGATCTCACTTGCTAACCAGGAGCTCCCAGGAT

>ZNF609|exon0-complete|874bp

CAATGATGTTGTCCACTGGGCATGTACTGACCAATGTGGCAGGTCTGAGAACATAGCTGAAGCTGAAAATAGGAAAGCTGGGGGCAAGGAAGAGCCTTGAATCTTGAGGTGGGACGTTGACTCTAAGATGTCCTTGAGCAGTGGAGCCTCCGGAGGGAAAGGAGTGGATGCAAACCCGGTTGAGACATACGACAGTGGGGATGAATGGGACATTGGAGTAGGGAATCTCATCATTGACCTGGACGCCGATCTGGAAAAGGACCAGCAGAAACTGGAAATGTCAGGCTCAAAGGAGGTGGGGATACCGGCTCCCAATGCTGTGGCCACACTACCAGACAACATCAAGTTTGTGACCCCAGTGCCAGGTCCTCAAGGGAAGGAAGGCAAATCAAAATCCAAAAGGAGTAAGAGTGGCAAAGACACTAGCAAACCCACTCCAGGGACTTCCCTGTTCACTCCAAGTGAGGGGGCAGCTAGCAAGAAAGAGGTGCAGGGGCGCTCAGGAGATGGTGCCAATGCTGGAGGCCTGGTTGCTGCTATTGCTCCCAAGGGCTCAGAGAAGGCGGCTAAGGCATCCCGCAGTGTAGCCGGTTCCAAAAAGGAGAAGGAGAACAGCTCATCTAAGAGCAAGAAGGAGAGAAGCGAAGGAGTGGGGACTTGTTCAGAAAAGGATCCTGGGGTCCTCCAGCCAGTTCCCTTGGGAGGACGGGGTGGTCAGTATGATGGAAGTGCAGGGGTGGATACAGGAGCTGTGGAGCCACTTGGGAGTATAGCTATTGAGCCTGGGGCAGCGCTCAATCCTTTGGGAACTAAACCGGAGCCAGAGGAAGGGGAGAATGAGTGTCGCCTGCTAAAGAAAGTCAAGTCTGAAAAG

***The Sanger-determined PCR product sequences of the twelve selected circRNAs***

> ASPHD1 – sanger sequencing result

CCGTACTTGACGACGTGCTCCCTTCACACACCCAGGCTGGAGAGACACTGCGCTCAGGGACGGCTTGATGGGAGAGGCTGGGGCAGGGCCTTATGTCTTCTCCTAGTACAACCAAACAGGCTCCCCCGAAGATGGGCCTCGAGTGGTCTTCATCGTGGACCTCTGGCACCCA

>ASPHD1 - reverse complement of sanger sequencing result

TGGGTGCCAGAGGTCCACGATGAAGACCACTCGAGGCCCATCTTCGGGGGAGCCTGTTTGGTTGTACTAGGAGAAGACATAAGGCCCTGCCCCAGCCTCTCCCATCAAGCCGTCCCTGAGCGCAGTGTCTCTCCAGCCTGGGTGTGTGAAGGGAGCACGTCGTCAAGTACGG

>CDYL – sanger sequencing result

TCTTCCCTTTTTTATTTTTCCTTTTGTCAACAATCCTTTCAACCTTTCCCGTTAACAGCTAAGCCTGTGGCCATGGCTGCAGTCACAGGGCCGGGCACCTGAGGCACTAGTGGGTGTATCCTGGGCCTAAGG

>CDYL – reverse complement of sanger sequencing result

CCTTAGGCCCAGGATACACCCACTAGTGCCTCAGGTGCCCGGCCCTGTGACTGCAGCCATGGCCACAGGCTTAGCTGTTAACGGGAAAGGTTGAAAGGATTGTTGACAAAAGGAAAAATAAAAAAGGGAAGA

>CCDC9 – sanger sequencing result

GGGCTGTGACATCCAGTGCGGACTTCCTGGACGACCCCCGGCGACGCAGCGGGCCCCTGGAGGAGTCTGAGCGGGACCGC

CGGGAGGAGAGCCGCCGGCACGGCCGCAACTGGGGGGGCCCCGACTTCGAGCGGGTGCGCTGTGGCCTTGAGCACGAGCG

GCAGGAGTGGGAGGAGCGGCGCAGGCAGAACAAA

>CCDC9 – reverse complement of sanger sequencing result

TTTGTTCTGCCTGCGCCGCTCCTCCCACTCCTGCCGCTCGTGCTCAAGGCCACAGCGCACCCGCTCGAAGTCGGGGCCCCCCCAGTTGCGGCCGTGCCGGCGGCTCTCCTCCCGGCGGTCCCGCTCAGACTCCTCCAGGGGCCCGCTGCGTCGCCGGGGGTCGTCCAGGAAGTCCGCACTGGATGTCACAGCCC

>DUS3L - sanger sequencing result

AAAATTGGTTTCTGGTGGGGGCTCCGGGCAGCAGGGGTGAGGGGTCTTGGGTCAGCCCACGTCCTGCCCTCCCTCCACGTAGCGGCCCTGGACGTGTGCCTCGGTTTCCCCATCTGGGCAGGGAGGCACTGACCCAGCCGGCGATGTCTCGGTGGAGGGTCAA

>DUS3L - reverse complement of sanger sequencing result

TTGACCCTCCACCGAGACATCGCCGGCTGGGTCAGTGCCTCCCTGCCCAGATGGGGAAACCGAGGCACACGTCCAGGGCCGCTACGTGGAGGGAGGGCAGGACGTGGGCTGACCCAAGACCCCTCACCCCTGCTGCCCGGAGCCCCCACCAGAAACCAATTTT

>GSE1 - sanger sequencing result

TGCCCGCCACTCGCAACAGGCGGAGGAGCCCAGAGGCATGAGCCATGAGCCCAAGTCCCCTTCGCTAGGGATGCTTTCCACCGCGACCAGGACAGTC

>intergenic – sanger sequencing result

CGTCGCATCCGGGTAGCGCCACTCTGCCCATTTCACCCGGTGCCCATCAAAGCCCCAGCGTCCGTCATTCAGCTGCTGCTGATCTCCTTCCAGGGCTTCCCAATTGC

>intergenic - reverse complement of sanger sequencing result

GCAATTGGGAAGCCCTGGAAGGAGATCAGCAGCAGCTGAATGACGGACGCTGGGGCTTTGATGGGCACCGGGTGAAATGGGCAGAGTGGCGCTACCCGGATGCGACG

>GLIS3 - sanger sequencing result

ACCCCATTCAATCCCAGGCCCTGTGGGGTTCCGATGTCCGGTGGAGACTCATGCTGCATGATCTTCCATTCATTCTGAAAAACCTGTGGCCAAGACGGTCAAATATCCAATGTCACTAATGACTCCTTTCAGGCAAAGTCCAATAAGTTATCCATGGTGTGGGTTATAAGCCTTTCCCAGGATTTGGAGGAAAAGGGCTTCCAAACTCCTGCTGCTTTGGCTTTAAAGTATGTGACCCTGACATGCCTCCAGCCTGGGTGACCTGGAATCGCACA

> NBPF14 - sanger sequencing result

CAAGCCCGCACTTCTGTAGGGCTGGCATGGGTCAGTCAGTTCAAGATAACCTGAAGGAGTCGAATAACATCTATCCAGTGAGTCCTGCAAGACTTCAGGCTCTTTCTCATCCAGCAGCTCCCTGCTGAGCCTGGGGCATGATGGGTCTTGGTCTTCTTCCGA

>NBPF14 - reverse complement of sanger sequencing result

TCGGAAGAAGACCAAGACCCATCATGCCCCAGGCTCAGCAGGGAGCTGCTGGATGAGAAAGAGCCTGAAGTCTTGCAGGACTCACTGGATAGATGTTATTCGACTCCTTCAGGTTATCTTGAACTGACTGACCCATGCCAGCCCTACAGAAGTGCGGGCTTG

>PROSC – sanger sequencing result

GCCCTGCCATTTTAGTTCGGACTGCTAGAAAAGCATCAAATCCCAAAATTCTGTCTTTGTGTCCTGAGATCAAATGGCACTTCATTGGCCACCTACAGAAACAAAATGTCAACAAATTGATGGGATCTCCCAGCCATCCAGCCCCGGCTAGTGGCGGTCAGCAAAACCAAACCTGCAGACATGGTGATCGAGGCCAATA

>RP11-196G11.1 - sanger sequencing result

AGAAGAAATAGAGCAGGAAGATAGGATCCAGGCCAGGTAGACAGAACCAGCGAGAGACACCAGGGAGCTCAGCAGCATCAGGACAGAGGCCCAGCGTGTCCGCAGGCAACCCTTCCCAGCTCCAGAGAAGGCAACACCGAGGGAGGCCCAGCACCACAGTCCATGGCAGACACATGGTTCAGACTTGGCTGATTGATCTAAGAAACTTTATTGCTCAGAACCTTCCCTCAA

>TSN4 – sanger sequencing result

AAGCTAAGGGGGTGGTGACTCTGCAGTTTACCTCTGGTCTTCAACCAGCCTCACTGACATCCCAGAGATCTCACTTGCTAACCAGGAGCTCCCAGGATCCAAGGCTTCAGATTCCTCCTTCTTTCTCATTGACATGGTGCTCTGGGCCAGCTCAGCCTGGGAGCCCCCAGTCCCTGGGGGAAGCAGCTGGAAGGTGGGGTCCAGTTCCAGGATCATCTGATTGAGGCTCTCCAGTGAGAAGTCAATGTAGGAGTCAAGGTCCTCTGGGG

>ZNF609 - sanger sequencing result

AGGGGAGGGAAATGAGTGTCGCCTGCTAAGAAGTCAAGTCTGAAAAGCAATGATGTTGTCCACTGGGCATGTACTGACCAATGTGGCAGGTCTGAGAACATAGCTGAAGCTGAAAATAGGAAAGCTGGGGGCAAGGAAGAGCCTTGAATCTTGAGGTGGGACGTTGACTCTAAGATGTCCTTGAGCAGTGGAGCCTCCGGAGGGAAAGGAGTGGATGCAAACCCGGTTGAGACATACGACAGTGGGGATGAATGGGACATTGGAGTAGGGAATCTCATCATTGACCTGGACGCCGATCTGGAAAAGGACCAGCAGAAACTGGAAATGTCAGGCTCA
